# A Chicken Tapasin ortholog can chaperone empty HLA molecules independently of other peptide-loading components

**DOI:** 10.1101/2023.06.23.546255

**Authors:** Georgia F. Papadaki, Claire H. Woodward, Michael C. Young, Trenton J. Winters, George M. Burslem, Nikolaos G. Sgourakis

## Abstract

Human Tapasin (hTapasin) is the main chaperone of MHC-I molecules, enabling peptide loading and antigen repertoire optimization across HLA allotypes. However, it is restricted to the endoplasmic reticulum (ER) lumen as part of the protein loading complex (PLC) and therefore is highly unstable when expressed in recombinant form. Additional stabilizing co-factors such as ERp57 are required to catalyze peptide exchange *in vitro*, limiting uses for the generation of pMHC-I molecules of desired antigen specificities. Here, we show that the chicken Tapasin (chTapasin) ortholog can be expressed recombinantly at high yields in stable form, independently of co-chaperones. chTapasin can bind the human HLA-B^*^37:01 with low micromolar-range affinity to form a stable tertiary complex. Biophysical characterization by methyl-based NMR methods reveals that chTapasin recognizes a conserved β_2_m epitope on HLA-B^*^37:01, consistent with previously solved X-ray structures of hTapasin. Finally, we provide evidence that the B^*^37:01/chTapasin complex is peptide-receptive and can be dissociated upon binding of high-affinity peptides. Our results highlight the use of chTapasin as a stable scaffold for future protein engineering applications aiming to expand the ligand exchange function on human MHC-I and MHC-like molecules.

## Introduction

The display of peptide-Major Histocompatibility Complex class I (pMHC-I) molecules on the cell surface is crucial for T cell and natural killer (NK) cell immunity. The loading of peptides onto MHC-I within cells relies heavily on the activity of the Protein Loading Complex (PLC), which comprises of several molecules including the transporter associated with antigen processing (TAP)1/2, Tapasin, calreticulin, ERp57, and the MHC-I-β_2_m heterodimer (1, 2). The intracellular enzymes endoplasmic reticulum aminopeptidase 1 and 2 (ERAP1/2) optimize the peptide cargos for MHC-I presentation, playing an indirect role in the modulation of adaptive immune responses (3–5). Tapasin is a transmembrane glycoprotein that acts as a bridge between the TAP transporter and MHC-I to facilitate peptide loading. Its significance in antigen presentation is evident in Tapasin-depleted cells by the reduced levels of surface MHC-I, which affects antigen presentation, CD8+ T cell development, and anti-viral immunity (6, 7). Additionally, many tumors evade immune surveillance by targeting components of the PLC, including Tapasin. The available crystal structures of Tapasin-ERp57 complex (8) and a high-resolution structure of Tapasin bound to B^*^44:05 (9) have proposed a mechanistic model where Tapasin interacts with the peptide binding domain of MHC-I via the α_2-1_ helix and the β-sheet floor, while the membrane proximal domain re-orients the MHC-I via the α_3_ domain and β_2_m (9). However, understanding in depth the mechanism by which Tapasin performs its peptide loading function has been challenging due to the dynamic flexibility of both the chaperone and the MHC-I molecule. Studies focusing on the Tapasin homolog, TAPBPR, which is not a component of the PLC have provided insights into Tapasin function (10–12). TAPBPR functions both as chaperone and a peptide-editor with distinct allelic specificity (13). We have recently characterized a chicken ortholog of TAPBPR (chTAPBPR) that interacts with a broader panel of HLAs (Human Leukocyte Antigens; the human MHCs) and has a strong preference for empty HLA-I molecules (14). However, further exploration of Tapasin/MHC-I interactions is necessary to understand chaperone dependence among allelic variations in HLA molecules.

In this work, we aim to expand the function and applications of chaperones to classical, non-classical, and MHC-like molecules. We characterize an ortholog of Tapasin from chicken (*Gallus gallus*) that has co-evolved with MHC-I to effectively promote peptide-loading (15), and show that it can be stably expressed in recombinant form without the need of co-chaperones. We further identify xeno reactivity with a panel of HLA allotypes using single antigen beads (SABs) and define low micromolar interactions with the disease-relevant HLA-B^*^37:01 allotype (16, 17). Using biophysical assays and solution NMR, we show that a stabilized B^*^37:01/chTapasin complex is peptide-receptive. Our results highlight the use of chicken Tapasin as a standalone molecular chaperone to stabilize empty, receptive HLA molecules, towards a range of *in vitro* and *in cell* applications (18–22).

## Results

To characterize differences between human Tapasin (hTapasin) and its ortholog from chicken (chTapasin), we expressed both proteins recombinantly in insect cells and purified them by size exclusion chromatography (SEC) (**Figure S1A**) (23). The chTapasin showed enhanced thermal stability with a melting temperature (T_m_) of 57.4 °C compared to ∼40 °C for hTapasin, as measured by differential scanning fluorimetry (DSF) (**Figure 1A, Figure S1B**). To thoroughly evaluate interactions with HLAs in a high-throughput manner, we created multivalent chaperone tetramers (24) and incubated them with single antigen beads (SABs), as previously described (14, 25). The beads are coated with 97 different HLA-I molecules that are loaded with a variety of peptides derived from EBV-transfected cell lines (26, 27). Using the monoclonal, pan-HLA class I antibody W6/32, we demonstrated comparable mean fluorescence intensity (MFI) levels for all allotypes (**Figure S2A**). As a negative control, we used a construct of the homologous chaperone TAPBPR carrying substitutions on important residues that have been shown to disrupt interactions with HLA-A^*^02:01 (hTAPBPR^TN6^: E205L, R207E, Q209S, and Q272S) (23, 28). As expected, staining with hTapasin tetramers revealed low MFI levels of binding, similar to hTAPBPR^TN6^, indicating the lack of interactions across the tested HLA panel in the absence of co-chaperones. Contrastingly, chTapasin showed a systematic shift towards higher MFI levels compared to both hTAPBPR^TN6^ and hTapasin, and a clear interaction with HLA-B^*^37:01 (**Figure 1B, Figure S2B-C**).

**Figure 1.**
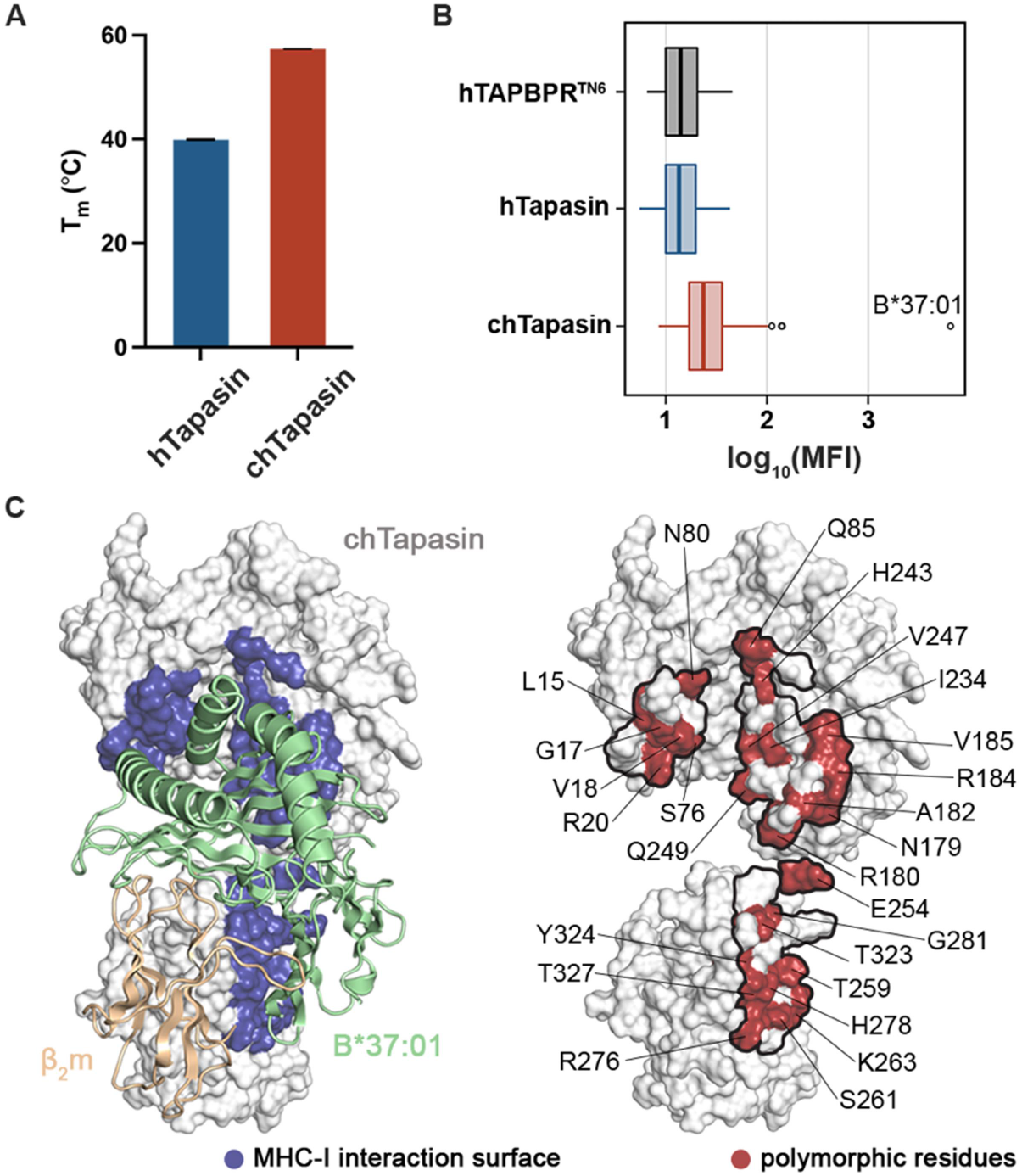
The chicken Tapasin ortholog shows enhanced stability and distinct interactions with HLA allotypes. **A**. Comparison of the thermal stabilities of human *vs*. chicken Tapasin by differential scanning fluorimetry (DSF) experiments. *T*_m_, melting temperature in degrees Celsius. The plotted data represent triplicate assays (*n* = 3) **B**. Box plot of the distribution of human and chicken Tapasin MFI levels for 97 different MHC-I allotypes. Human TAPBPR carrying the substitutions E205L, R207E, Q209S, and Q272S (hTAPBPR^TN6^) was used as negative control. The left boundary of the box represents the 25th percentile, the line within the box represents the median, and the right boundary of the box represents the 75th percentile. Whiskers extending above and below the box indicate the 10th and 90th percentiles, respectively. Any points appearing above the whiskers represent outliers that fall beyond the 90th percentile. Plotted data are mean from three replicates (*n* = 3). **C**. Surface representation of the chTapasin bound to HLA-B*37:01 structure model generated using the BAKER-ROBETTA server (29). The predicted contact surfaces of chTapasin with the heavy chain of B*37:01 are highlighted in blue (left). Surface representation of chTapasin where the polymorphic residues within the contact surfaces (denoted by the black line) are marked in red (right).

We next examined to what extent these HLA interaction trends can be determined by sequence divergence between human and chicken Tapasin. For this purpose, we generated a model of chTapasin in complex with HLA-B^*^37:01 using the BAKER-Robetta server (29), based on the crystal structure of hTapasin with HLA-B^*^44:05^T73C^ (PDB ID:7TUE) (9). Comparison of the interacting residues with the MHC-I heavy chain within 3.5 Å (**Figure 1C**) revealed that the majority is polymorphic between the two orthologs and this variation could enable interactions with distinct HLA allotypes (**Figure 1C, Figure S3**). Interestingly, even though multiple chTapasin allotypes have been described (15), neither of the polymorphic sites are located within the interacting surfaces with MHC-I.

To thoroughly investigate the interaction between chTapasin and HLA-B^*^37:01, we used surface plasmon resonance experiments (SPR), where chTapasin was immobilized to the chip surface at ∼2000 RU. HLA-B^*^37:01 refolded with the conditional peptide FEDLRVJSF (photo B37; J = 3- amino-3-(2-nitrophenyl)-propionic acid) that gets cleaved upon UV irradiation (30, 31) was flown over the chip at different concentrations. No binding was observed with either peptide-loaded or -deficient wild-type (WT) B^*^37:01 molecules (**Figure S4A-B**). We hypothesized that the lack of interaction could be due to the instability of empty HLA-B^*^37:01 molecules. In a recently proposed universal platform for the generation of MHC-I molecules with enhanced stability, it was suggested to introduce an interchain disulfide bond between the light and heavy chains (32). Therefore, we used an engineered HLA-B^*^37:01 molecule, referred to as open B^*^37:01, that carried the substitutions H31C and G120C in the light and heavy chains, respectively (32). We found that chTapasin showed low micromolar affinity to the UV-irradiated (empty), open B^*^37:01 with a K_D_ of ∼1.2 μM (**Figure 2A**), but not with the open, loaded B^*^37:01 (**Figure S4C**). This is likely due to rapid light-chain dissociation from the WT empty MHC-I, as a result of the known cooperativity between the peptide and β_2_m for binding to the heavy chain (33, 34). Contrarily, the covalent attachment of the β_2_m promotes the open state of the peptide-binding groove, facilitating interactions with the chaperone (**Figure 2B**).

**Figure 2.**
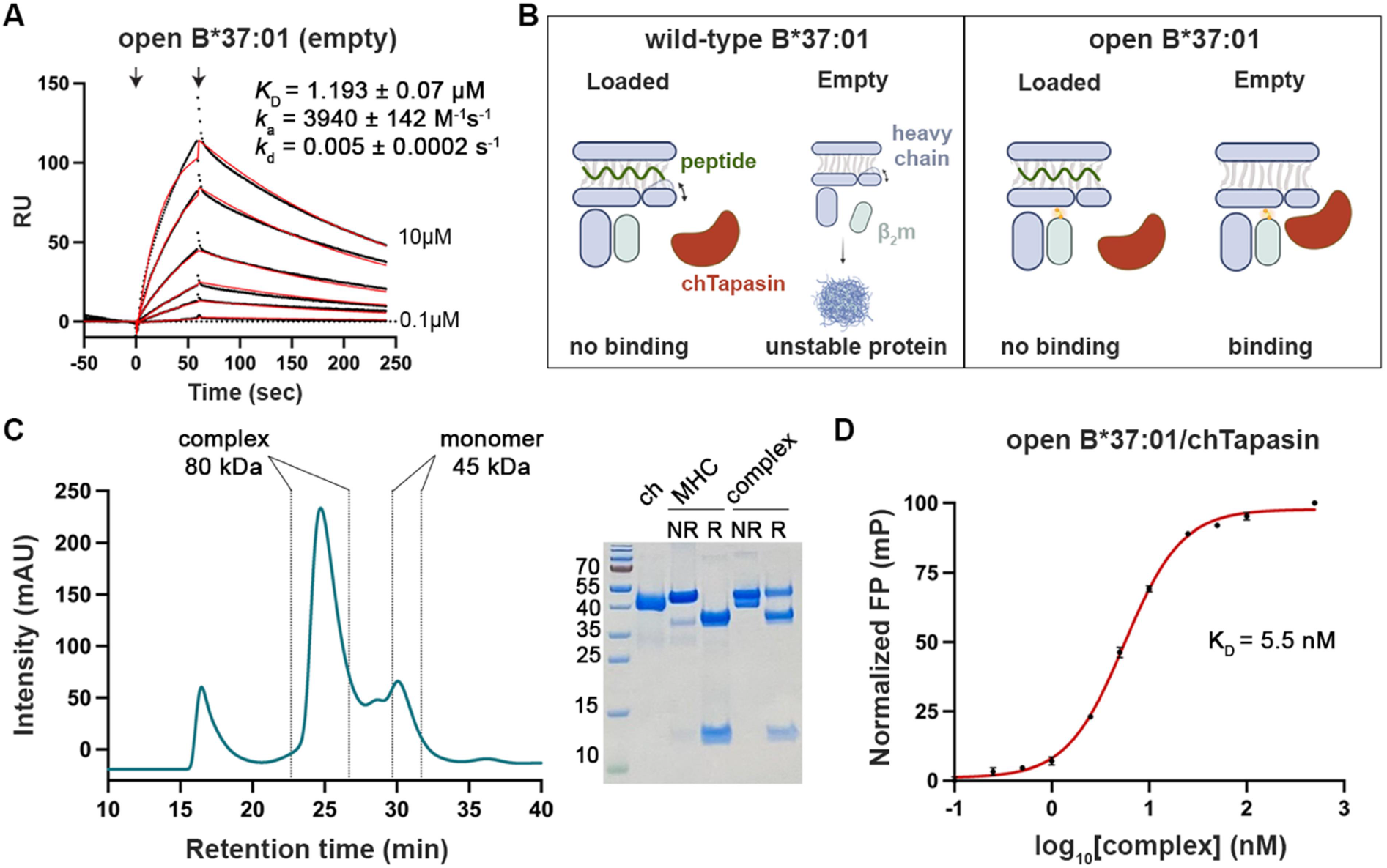
The chTapasin ortholog binds to the open, empty HLA-B*37:01 forming a stable peptide-receptive complex. **A**. Representative sensorgrams of UV-irradiated HLA-B*37:01. *K*_D_, equilibrium constant; *k*_a_, association rate constant; *k*_d_, dissociation rate constant; RU, resonance units. Fits from the kinetic analysis are shown in red lines and the concentrations **B**. Schematic representation of the chTapasin interactions with wild-type versus open HLA-B*37:01. Created with BioRender.com. **C**. SEC analysis of the mixture of chTapasin (ch) with open HLA-B*37:01/FEDLRVJSF (MHC) upon 40 mins UV-irradiation. The peak corresponding to the empty complex was collected and all the components were identified by SDS/PAGE under reduced (R) or non-reduced (NR) conditions. **D**. FP saturation binding curve of _FITC_KEDLRVSSF with increasing concentrations of open B*37:01/chTapasin complex. Log_10_(K_D_) = 0.74. Results of three replicates (mean ± SD) are plotted.

To explore further whether the open B^*^37:01/chTapasin was receptive for incoming peptides, we purified the empty complex by incubating 1:1.3 stoichiometric ratio of chTapasin and UV-irradiated open B*37:01, followed by SEC (**Figure 2C**). Next, we followed the binding of a fluorescently labeled, high-affinity peptide by real-time fluorescence polarization (FP) (35). Using a series of complex concentrations and an optimized concentration of the fluorescently labeled peptide (_FITC_KEDLRVSSF) (32), we observed improved binding of the FITC-peptide at higher concentrations (**Figure S5**). After 1 h, FP values reached a maximum plateau with an estimated affinity (K_D_) at the low nanomolar range (K_D_ = 5.5 nM) (**Figure 2D**) (35, 36), suggesting that the open B*37:01/chTapasin is peptide-receptive.

To evaluate the effects of chTapasin binding to the open B^*^37:01 complex in a solution environment by NMR, we employed established selective isotopic labeling methods for MHC-I molecules (37, 38). We focused on the alanine, isoleucine, leucine, and valine side chain methyl groups in β_2_m, which are known to be impacted by Tapasin binding (9), as highly sensitive probes for changes in the local magnetic environment. We first assigned the 2D methyl HMQC spectrum for the light chain of open B^*^37:01 loaded with the photo B37 peptide (**Figure 3A**). We then incubated the open MHC-I with excess of chTapasin and UV-irradiated the peptide to generate the empty complex. Overlay of the resulting 2D methyl spectra with the apo-open B^*^37:01/photo B37 spectra shows significant chemical shift effects upon Tapasin binding, where the interaction between the two proteins is slow on the NMR timescale (**Figure 3A-D**). We observed that several methyl probes appear as two peaks in the chTapasin-bound NMR spectra, corresponding to the apo-open B^*^37:01 state and the chTapasin-bound state. Specifically, residues Val9 and Val93 are located on the β-sheet (residues 6-10) and the C-terminus of β_2_m that interact with chTapasin as previously reported for hTapasin (9). The methyl groups with affected 2D HMQC resonances corresponding to residues Leu23, Val27, Val37, Leu40, and Val82 are dispersed throughout the β-sheets of the light chain, indicating widespread allosteric effects on the β_2_m structure induced by binding to chTapasin and peptide release (**Figure 3E**). Additionally, the observed chemical shift perturbations for residues Ile35, Val85, and Leu87 that are located beneath the peptide binding groove, supporting previous data that indicate a possible mechanism for peptide release by interfering with the β_2_m/α_2_ interface of the MHC (9). These results are consistent with a prolonged lifetime of the open B^*^37:01/chTapasin protein complex (>200 msec timescale), which exists as an equilibrium of unbound and bound states in solution.

**Figure 3.**
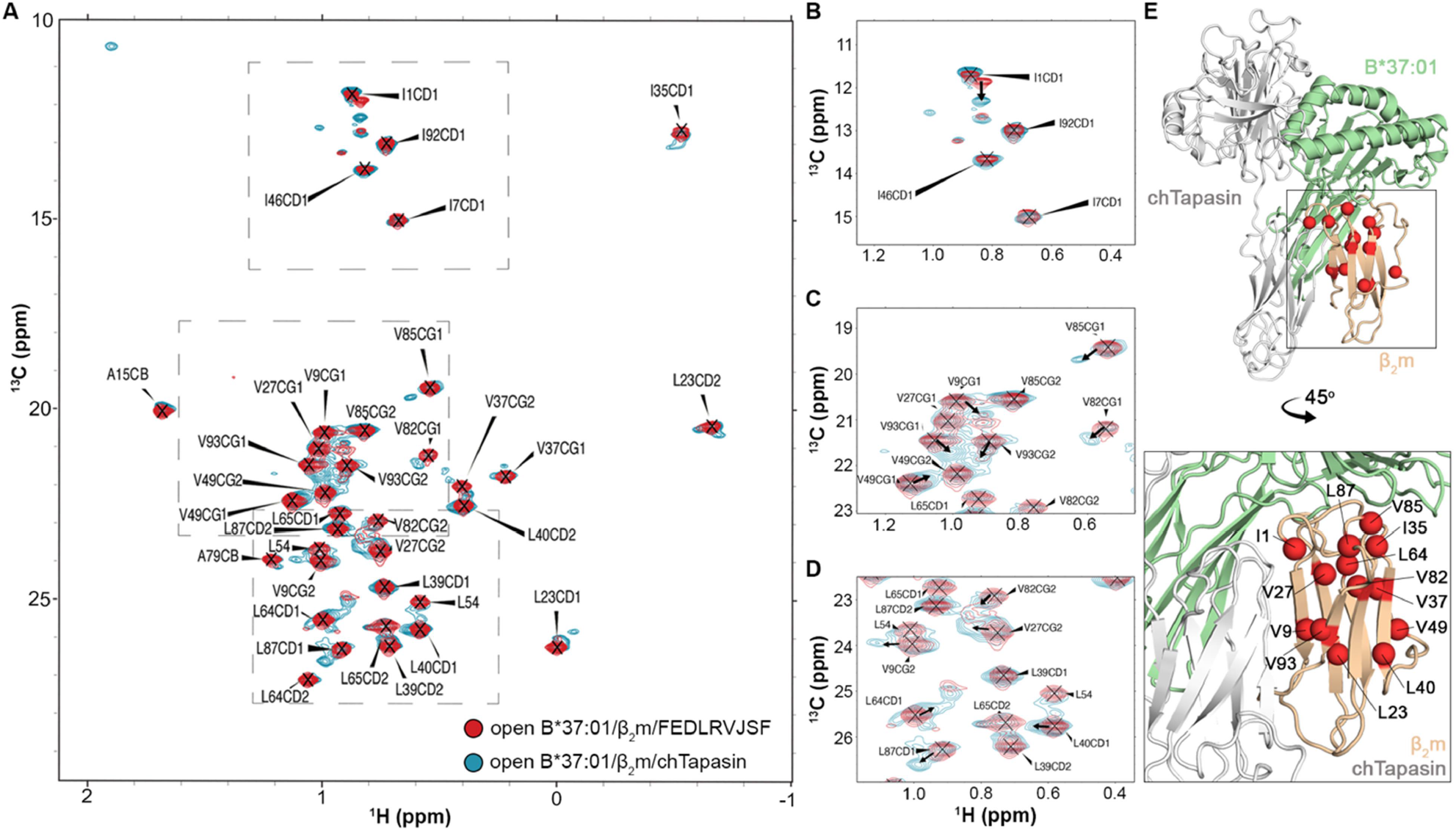
Conformational changes in open B*37:01/β_2_m induced by interaction with chTapasin. **A**. 2D ^1^H-^13^C SOFAST HMQC spectra of U-[^15^N, ^2^H], ^13^CH_3-_labeled alanine, isoleucine, leucine, and valine residues of human β_2_m (H31C) in the B*37:01 (G120C)-FEDLRVJSF bound state (in red). The sample was subsequently incubated with chicken Tapasin and UV irradiated for 40 min prior to collecting additional ^1^H-^13^C SOFAST HMQC spectra (in cyan). Both spectra were collected at 600 MHz ^1^H magnetic field, at 25°C. Due to lack of stereospecific assignments, the CD1 and CD2 methyl groups of Leu54 are not indicated. **B-D**. Zoomed regions of the ^1^H-^13^C SOFAST HMQC spectra with arrows highlighting methyl probes experiencing minor chemical shift changes upon chTapasin binding, indicated by a slow-exchanging minor state peak. **E**. Methyl probes that show a detectable slow-exchanging minor peak upon chTapasin binding are shown as red spheres on the predicted B*37:01/β_2_m/chTapasin model by BAKER-ROBETTA (29).

## Discussion

Cells detect foreign or abnormal antigens displayed by MHC-I molecules via T cell receptors (TCRs), initiating the first crucial stage in the development of immunity against viruses and tumors (39). Considering the fundamental importance of T cell responses, fast, high-throughput, and cost-effective methods, including peptide exchange technologies for pMHC-multimers generation, are emerging to study pMHC:TCR antigen recognition. The molecular chaperones Tapasin and TAPBPR determine to a high extent the resulting peptide repertoire (19, 23, 28, 40–42). We have previously described a robust method to prepare biologically relevant, stable pMHC-I molecules loaded with peptides of choice by exploiting hTAPBPR (20) that can function independently of the PLC, albeit this approach is limited to specific allotypes that can be recognized by the chaperone (25). Aiming to expand the HLA-I repertoire susceptible to TAPBPR-assisted peptide exchange, we characterized a TAPBPR ortholog from chicken (14). While the predicted structure of chTAPBPR exhibits remarkable similarity to hTAPBPR (13), their sequence variability conferred an expanded interaction pattern to include representatives from six additional supertypes (14, 43, 44).

Human Tapasin is an indispensable part of the PLC and serves as the primary chaperone for MHC-I molecules, facilitating the loading of peptides and optimizing the antigen repertoire across various allotypes (45). However, Tapasin expression is restricted to the PLC, and therefore is highly unstable when produced in recombinant form. To address this limitation, we explore chicken Tapasin, which is encoded by diverse protein sequences, each closely associated with the expression of the dominant MHC molecules (15). We show that chTapasin functions without co-chaperones and interacts tightly with an engineered, stabilized HLA-B^*^37:01 (32), suggesting that variation of the amino acid sequence between Tapasin orthologs is accountable for distinct MHC-I interactions. Importantly, deep mutational scanning on hTAPBPR revealed two prominent regions that drive these interactions, which with minimal alterations allow for peptide exchange on desired HLA allotypes (14). In an analogous manner, performing a single site-saturation mutagenesis (SSM) library on hTapasin (14), would provide insights on important residues that could be substituted to guide the HLA allelic specificity of hTapasin. Notably, HLA-B^*^37:01 is a low hTapasin-dependent allotype (45) that has been associated with diseases like multiple sclerosis and flu (16, 17), underscoring the importance of uncovering new tools to expand chaperone interactions. Overall, our results highlight the potential use of chTapasin as an orthologous chaperone, and provide a readily available framework for future protein engineering applications aimed at broadening the ligand exchange function of human MHC I and MHC-like molecules.

## Experimental Procedures

### Sequence analysis

The sequences used in this study are: hTAPBPR (Q9BX59), chTAPBPR (NP_001382952.1), hTapasin (O15533), and chTapasin (A4F5A9). Alignments were performed in ClustalOmega (46) and processed in ESPript (47).

### Recombinant protein expression, refolding, and purification

The luminal domains of human and chicken Tapasin, and hTAPBPR^TN6^ encoding the BirA substrate peptide (BSP; LHHILDAQKMVWNHR) and 6-His tag were stably expressed in the *Drosophila melanogaster* S2 cells and purified as previously described (23).

The wild-type MHC-I heavy chain tagged with BSP and the β_2_m light chain were provided by the NIH (Emory university) or synthesized and cloned into pET-22b(+) vector (Genscript). For the open constructs, substitutions in positions H31C (β2m) and G120C (heavy chain) were generated using site-directed mutagenesis (32). Upon *Escherichia coli* BL21 (DE3) transformation (New England Biolabs), expressed proteins were isolated from inclusion bodies. For pMHC-I generation, we performed *in vitro* refolding as previously described (30).

### Purification of empty MHC-I/Tapasin complex

chTapasin was incubated with open B^*^37:01/FEDLRVJSF at 1:1.3 molar ratio for 1 hour at room temperature (RT), followed by 40-min UV irradiation at 365 nm and 30 min additional RT incubation (14). The empty complex was purified by SEC using a Superdex 200 pg Increase 10/300 GL. To identify all components, the eluted peaks were analyzed by SDS-polyacrylamide gel electrophoresis (PAGE).

### Peptides

Peptide sequences are given as single-letter codes. Photolabile peptides were purchased from Biopeptek Inc. (Malvern, USA) using J as 3-amino-3-(2-nitrophenyl)-propionic acid. FITC-labeled peptides were synthesized as previously described (32). Peptides were solubilized in distilled water and centrifuged at 14,000 rpm for 15 min before measuring their concentration at 205 nm using the respective extinction coefficient.

### Differential scanning fluorimetry

The thermal stability of Tapasin was measured by incubating 7 μM of protein with 10X SYPRO Orange dye in PBS buffer in 20 μL. Samples were loaded in a MicroAmp Optical 384-well plate in triplicates and analyzed on a QuantiStudio 5 real-time PCR machine. Excitation and emission wavelengths were set to 470 and 569 nm, and temperature increased at a rate of 1°C/min between 25-95°C. Data analysis and fitting were performed in GraphPad Prism v9.

### Biotinylation and tetramer formation

BSP-tagged proteins were biotinylated using the BirA biotin-ligase bulk reaction kit (Avidity), according to the manufacturer’s instructions. Streptavidin-PE (Agilent Technologies, Inc.) at 4:1 monomer:streptavidin molar ratio was added in the dark, every 10 min at RT over 10-time intervals (20).

### Single antigen bead assay

Tetramerized Tapasin orthologs and hTAPBPR^TN6^ (7 μM) were mixed with 4 μL of LABScreen SABs (OneLambda Inc., CA, USA) and incubated for 1 hour, 550 rpm at RT. After four washes with the provided buffer to remove excess of tetramers, beads were resuspended in PBS buffer. The levels of peptide-loaded MHC-I on the beads were tested using the PE-conjugated W6/32 antibody (Biolegend, 311406). Binding levels were measured in Luminex 100 Liquid Analyzer System and the results analyzed in GraphPad Prism v9.

### Surface plasmon resonance

SPR experiments were performed in a BiaCore X100 instrument (Cytiva) in SPR buffer (150 mM NaCl, 20 mM sodium phosphate pH 7.4, 0.1% Tween-20). Approximately 2000 resonance units (RU) of biotinylated chTapasin were conjugated on a streptavidin-coated chip (Cytiva) at 10 μL/min. Various concentrations of pMHC-I were flown over the chip for 60 sec at 30 μL/min followed by a buffer wash with 180 sec dissociation time, at 25°C. SPR sensorgrams, association/dissociation rate constants (k_a_, k_d_), and equilibrium dissociation constant K_D_ values were analyzed in BiaCore X100 evaluation software (Cytiva) using kinetic analysis settings of 1:1 binding. SPR sensorgrams and affinity-fitted curves were prepared in GraphPad Prism v9.

### Fluorescence polarization

FP was used to monitor the kinetic association of fluorescently labeled peptides and MHC-I. Various concentrations of open B^*^37:01/chTapasin complex were incubated with an optimized concentration of _FITC_KEDLRVSSF (10 nM) in FP buffer (PBS and 0.05% Tween-20). Excitation and emission values were 475 and 525 nm, respectively.

### NMR spectroscopy

An NMR sample of open HLA-B^*^37:01/β_2_m/FEDLRVJSF was prepared by *in vitro* refolding, by selectively labeling the light chain hβ_2_m with a {U-[^15^N,^2^H]; Ala, Ile, Leu, Val-[^13^CH_3_]}-labeled isotope-selective labeling scheme using established protocols and reagents (11, 48). The purified sample was prepared in standard NMR buffer (150 mM NaCl, 20 mM sodium phosphate pH 7.2, 0.001 M sodium azide, 5% D_2_O) in the presence of 2-fold molar excess of peptide. 2D SOFAST methyl ^1^H-^13^C HMQC and 2D ^1^H-^15^N TROSY spectra were collected. Methyl resonance assignments for the light chain were obtained using 3D SOFAST Cm-CmHm NOESY experiments recorded with 150 ms mixing time (49). The complex was then incubated with chTapasin and UV-irradiated for 40 minutes to cleave the photo B37 peptide. Approximately 70 μM of sample was prepared in identical buffer conditions with excess of chTapasin to collect 2D SOFAST methyl ^1^H-^13^C HMQC and 2D ^1^H-^15^N TROSY spectra for the complex. All NMR data were recorded at 600 MHz and 298 K, processed with NMRPipe (50), and analyzed using POKY (51).

## Data availability

All other data are contained within this article and in the supporting information. Final methyl assignments were deposited in the Biological Magnetic Resonance Data Bank (http://www.bmrb.wisc.edu) under accession number 51999.

## Supporting information

This article contains supporting information.

## CRediT author statement

N.G.S. and G.F.P conceptualization; N.G.S. and G.F.P. investigation; M.C.Y, T.J.W. and G.M.B. resources; G.F.P. and C.H.W. formal analysis; G.F.P. visualization; N.G.S. and G.F.P writing–original draft; N.G.S., G.F.P., C.H.W., M.C.Y, T.J.W. and G.M.B. writing–review & editing; N.G.S. supervision; N.G.S. funding acquisition.

## Acknowledgments

We acknowledge Dr. Monos and the CHOP Immunogenetics Laboratory for discussions and access to equipment and reagents. Authors N.G.S and G.F.P. are co-inventors in provisional patent applications related to this work. This research was supported through grants by NIAID (5R01AI143997), NIGMS (5R35GM125034), and NIDDK (5U01DK112217) to N.G.S.

