## Supporting information for "A Chicken Tapasin ortholog can chaperone empty HLA molecules independently of other peptide-loading components"

**This PDF file includes:**

Figures S1 to S5

### Figures

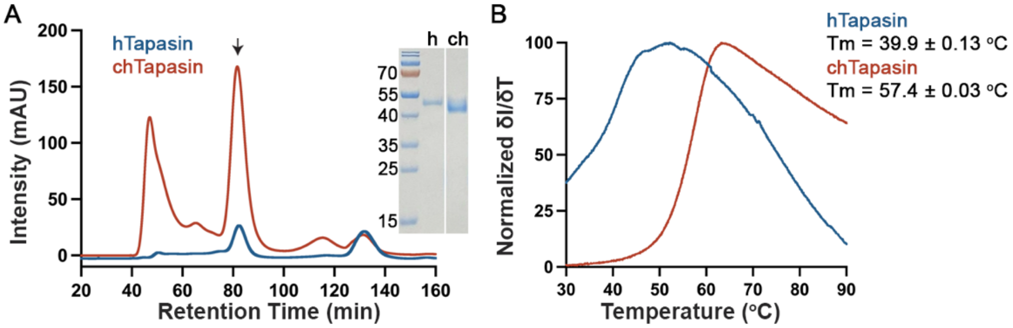


**Figure S1. Chicken Tapasin can be expressed in high yields with improved stability. A.** Size exclusion chromatography (SEC) traces of human (h) and chicken (ch) Tapasin proteins expressed in insect cells. The protein peaks are indicated by the arrow and were further validated by SDS-PAGE analysis. **B.** Differential scanning fluorimetry (DSF) of human versus chicken Tapasin. Melting temperatures are in degrees Celsius ($T_m$). Data are mean ± SD obtained from n = 3 technical replicates.

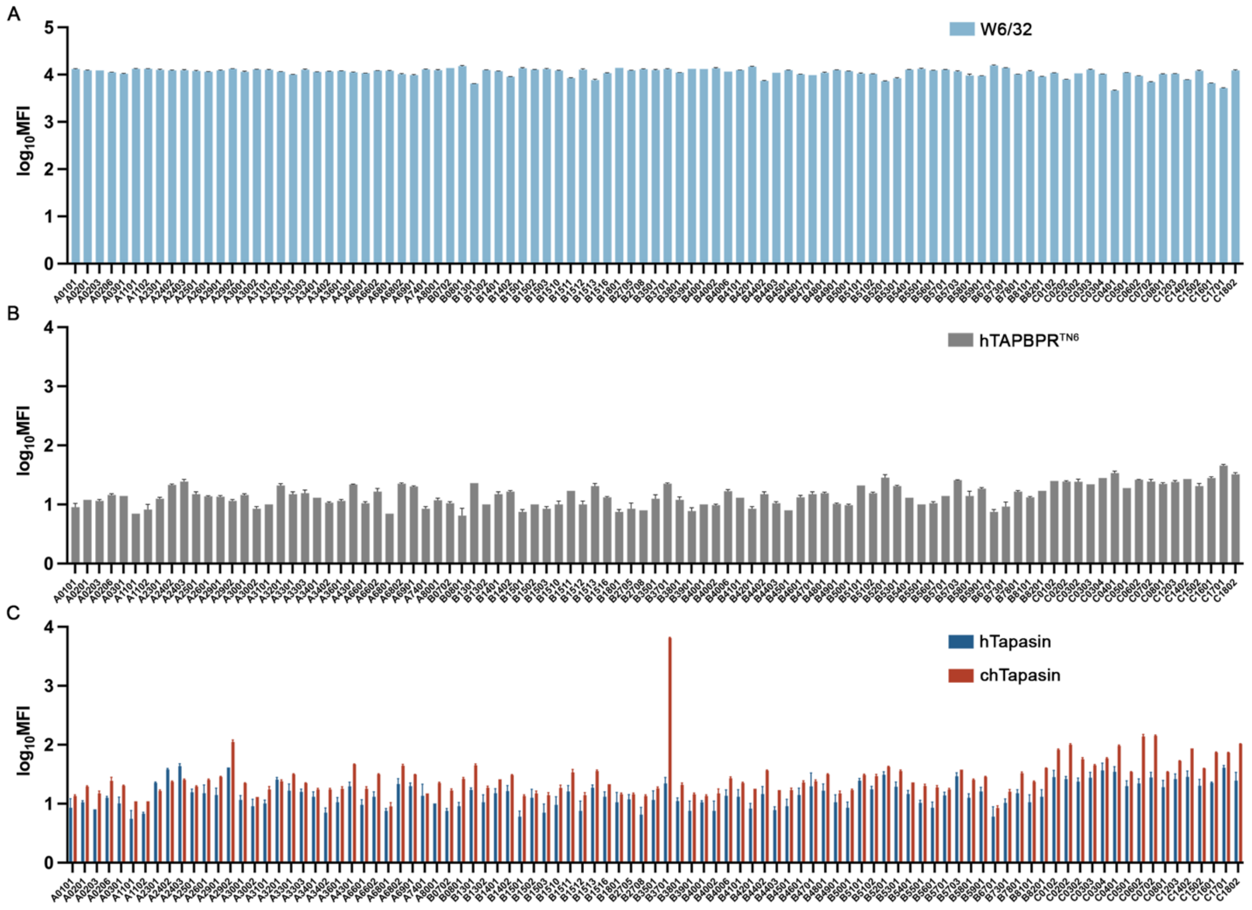


**Figure S2. Binding levels of human and chicken Tapasin on HLA single antigen beads. A.** Similar levels of peptide-loaded MHC-I molecules on the beads were observed upon staining with the primary anti-HLA class I antibody W6/32 conjugated with PE (Biolegend, 311406). Bar graphs showing the binding levels of tetramerized **B.** hTAPBPR$^{TN6}$ (negative control), and **C.** hTapasin and chTapasin to 97 different HLA allotypes on the SABs expressed as logarithm of Mean Fluorescence Intensity (MFI). The plotted data are mean ± SD of n = 3 independent experiments.

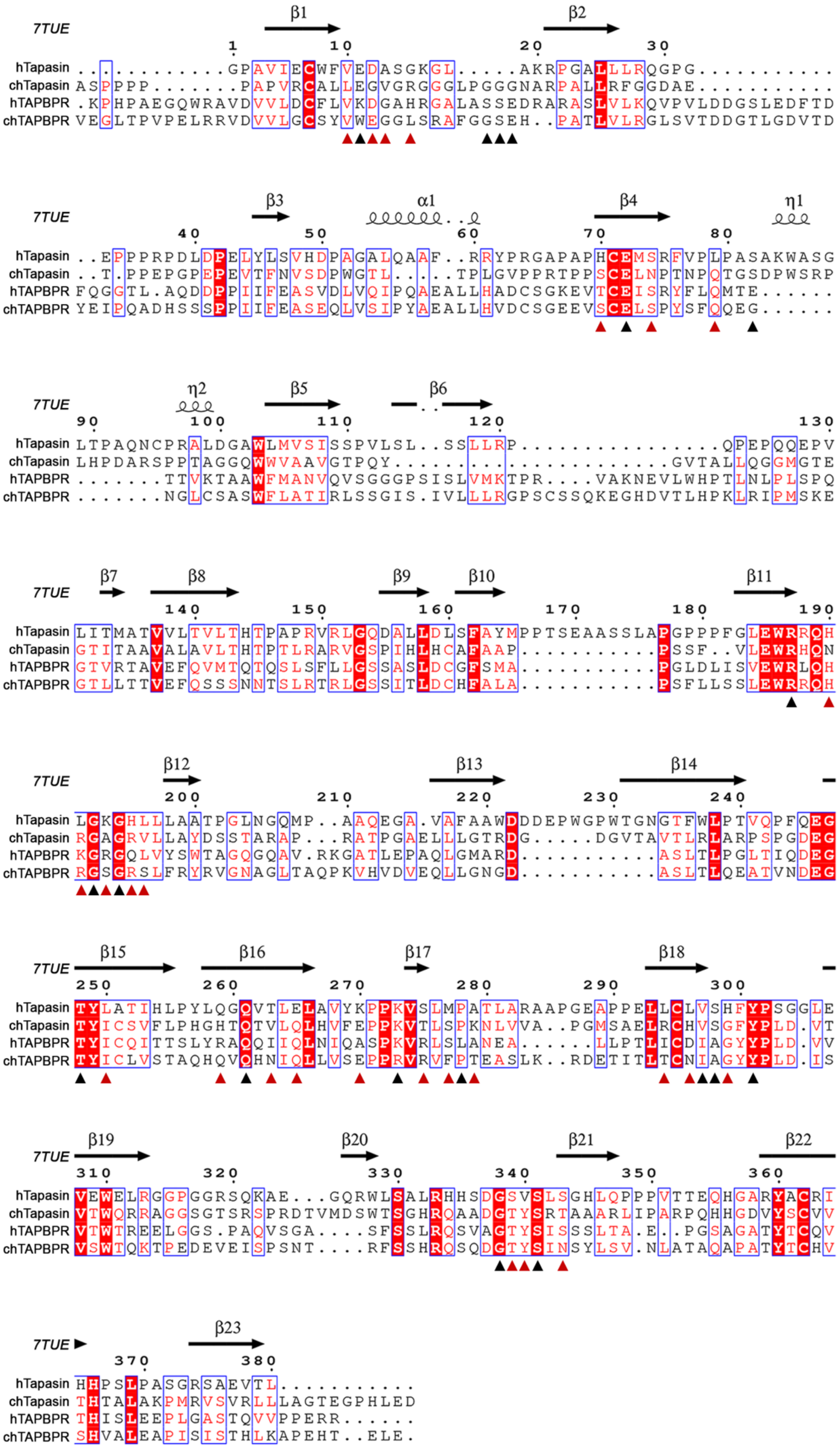
7TUE
β1 β2 β3 α1 β4 η1 η2 β5 β6 β7 β8 β9 β10 β11 β12 β13 β14 β15 β16 β17 β18 β19 β20 β21 β22 β23
hTapasin ...........GPAVIECWFVEDASGKGL.....AKRPGALLLRQGPG..........
chTapasin ASPPPP.......PAPVRCALLEGVGRGGLPGGGNARPALLRFGGDAE..........
hTAPBPR .KPHPAEGQWRAVDVVLDCFLVKDGAHRGALASSEDRARASLVLKQVPVLDDGSLEDFTD
chTAPBPR VEGLTPVPELRRVDVVLGCSYVWEGGLSRAFGGSEH..PATLVLRGLSVTDDGTLGDVTD
hTapasin ..EPPPRPDLDPELYLSVHDPAGALQAAF..RRYPRGAPAPHCEMSRFVPLPASAKWASG
chTapasin ..T.PPEPGPEPEVTFNVSDPWGTL....TPLGVPPRTPPSCELNPTNPQTGSDPWSRP
hTAPBPR FQGGTL.AQDDPPIIFEASVDLVQIPQAEALLHADCSGKEVTCEISRYFLQMTE......
chTAPBPR YEIPQADHSSSPPIIFEASEQLVSIPYAEALLHVDCSGEEVSCELSPYSFQQEG......
hTapasin LTPAQNCPRALDGAWLMVSISSPVLSL..SSLLRP...............QPEPQQEPV
chTapasin LHPDARSPPTAGGQWWVAAVGTPQY.......................GVTALLQGGMGTE
hTAPBPR .......TTVKTAAWFMANVQVSGGGPSISLVMKTPR....VAKNEVLWHPTLNLPLSPQ
chTAPBPR .......NGLCSASWFLATIRLSSGIS.IVLLLRGPSCSSQKEGHDVTLHPKLRIPMSKE
hTapasin LITMATVVLTVLTHTPAPRVRLGQDALLDLSFAYMPPTSEAASSLAPGPPPFGLEWRRQH
chTapasin GTITAAVALAVLTHTPTLRARVGSPIHLHCAFAAP...........PSSF..VLEWRHQN
hTAPBPR GTVRTAVEFQVMTQTQSLSFLLGSSASLDCGFSMA...........PGLDLISVEWRLQH
chTAPBPR GTLLTTVEFQSSSNNTSLRTRLGSSITLDCHFALA...........PSFLLSSLEWRRQH
hTapasin LGKGHLLLAATPGLNGQMP..AAQEGA.VAFAAWDDDEPWGPWTGNGTFWLPTVQPFQEG
chTapasin RGAGRVLLAYDSSTARAP...RATPGAELLLGTRDG.....DGVTAVTLRLARPSPGDEG
hTAPBPR KGRGQLVYSWTAGQGQAV.RKGATLEPAQLGMARD...........ASLTLPGLTIQDEG
chTAPBPR RGSGRSLFRYRVGNAGLTAQPKVHVDVEQLLGNGD...........ASLTLQEATVNDEG
hTapasin TYLATIHLPYLQGQVTLELAVYKPPKVSLMPATLARAAPGEAPPELLCLVSHFYPSGGLE
chTapasin TYICSVFLPHGHTQTVLQLHVFEPPKVTLSPKNLVVA..PGMSAELRCHVSGFYPLD.VT
hTAPBPR TYICQITTSLYRAQQIIQLNIQASPKVRLSLANEA......LLPTLICDIAGYYPLD.VV
chTAPBPR TYICLVSTAQHQVQHNIQLLVSEPPRVRVFPTEASLK..RDETITLTCNIAGYYPLD.IS
hTapasin VEWELRGGPGGRSQKAE...GQRWLSALRHHSDGSVSLSGHLQPPPVTTEQHGARYACRI
chTapasin VTWQRRAGGSGTSRSPRDTVMDSWTSGHRQAADGTYSRTAAARLIPARPQHHGDVYSCVV
hTAPBPR VTWTREELGGS.PAQVSGA....SFSSLRQSVAGTYSISSSLTA.E..PGSAGATYTCQV
chTAPBPR VSWTQKTPEDEVEISPSNT....RFSSHRQSQDGTYSINSYLSV.NLATAQAPATYTCHV
hTapasin HHPSLPASGRSAEVTL...........
chTapasin THTALAKPMRVSVRLLLAGTEGPHLED
hTAPBPR THISLEEPLGASTQVVPPERR......
chTAPBPR SHVALEAPISISTHLKAPEHT..ELE.

**Figure S3. Sequence alignment of human and chicken Tapasin and TAPBPR.** Alignment of the luminal domains of human Tapasin (hTapasin; UniProt: O15533), chicken Tapasin (chTapasin; UniProt: A4F5A9), human TAPBPR (hTAPBPR; UniProt: Q9BX59), and chicken TAPBPR (chTAPBPR; NP_001382952.1). Conserved or polymorphic residues between human and chicken Tapasin on the interface with HLA-B*37:01 are shown in black or red, respectively. The secondary structure of human Tapasin (PDB ID: 7TUE) is provided as reference (1). Conserved residues are marked in blue boxes. Alignment was performed in ClustalOmega (2) and processed in ESPript (3).

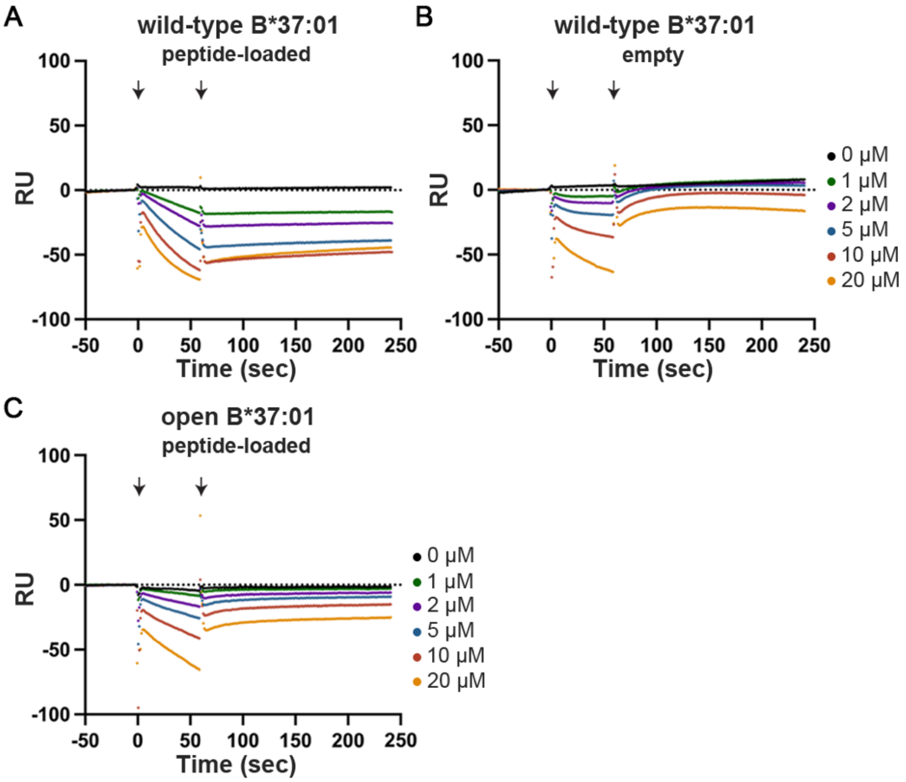


**Figure S4. Chicken Tapasin interactions are restricted to the open, empty B*37:01.** SPR sensorgrams of varying concentrations of soluble wild-type **A.** peptide-loaded, and **B.** -deficient B*37:01 or **C.** open, peptide-loaded B*37:01 flown over a streptavidin chip coupled with biotinylated chTapasin. Injection and washing start points are indicated by arrows. RU, resonance units.

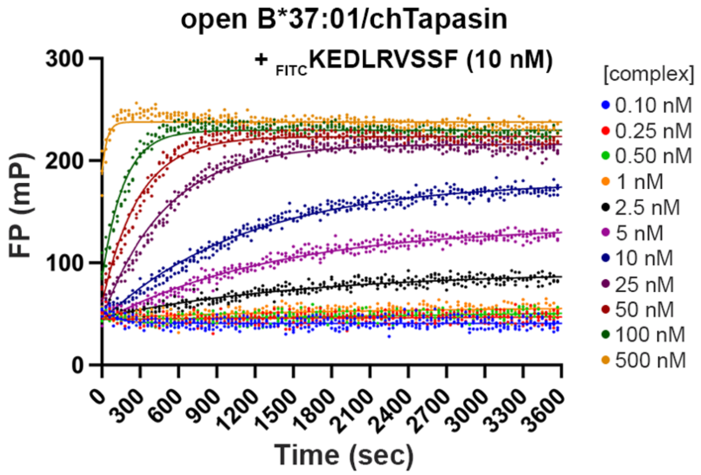


**Figure S5. The open B*37:01/chTapasin complex is peptide receptive.** Association profile of the fluorophore-conjugated peptide $_{\text{FITC}}$KEDLRVSSF (10 nM) to a series of open B*37:01/chTapasin complex concentrations. The data were fitted to a one-phase association model. Data from n = 3 technical replicates are plotted. FP, fluorescence polarization.
